# Fractal complexity of Escherichia coli nutrient transport channels is influenced by cell shape and growth environment

**DOI:** 10.1101/2023.11.29.569150

**Authors:** Beatrice Bottura, Liam Rooney, Morgan Feeney, Paul A. Hoskisson, Gail McConnell

## Abstract

Recent mesoscopic characterisation of nutrient-transporting channels in *E. coli* has allowed the identification and measurement of individual channels in whole mature biofilms. However, their complexity under different physiological and environmental conditions remains unknown. Analysis of confocal micrographs of biofilms formed by cell shape mutants of *E. coli* shows that channels have a high fractal complexity, regardless of cell phenotype or growth medium. In particular, biofilms formed by the mutant strain Δ*ompR*, which has a wide-cell phenotype, have a higher fractal dimension when grown on rich medium than when grown on minimal medium, with channel complexity affected by glucose and agar concentration in the medium. Osmotic stress leads to a dramatic reduction in Δ*ompR* cell size, but has a limited effect on channel morphology. This work shows that fractal image analysis is a powerful tool to quantify the effect of phenotypic mutations and growth environment on the morphological complexity of internal *E. coli* biofilm structures. If applied to a wider range of mutant strains, this approach could help elucidate the genetic determinants of channel formation in *E. coli* biofilms.

## 1. Introduction

The formation of spatial patterns is ubiquitous in biological systems, where discrete entities come together to form complex structures in a process called ‘‘morphogenesis’’ [1]. Bacterial populations are no exception to this phenomenon: bacterial biofilms exhibit a variety of internal patterns, from surface and 3D features to complex fractal shapes [2]. Fractal geometry has previously been proposed as a tool for the investigation of microbial growth patterns [3], and it has since been employed to quantify biofilm morphology from microscopy images [4]–[6], to describe colony morphogenesis in Gram-negative rod-shaped bacteria [7] and to analyse nutrient-limited growth patterns [5], [8].

In *E. coli*, fractal patterns mostly exist within cellular aggregates of co-cultured isogenic strains expressing different fluorescent makers [9] [10]. Fractal boundaries are formed during uniaxial cell growth and division, and can be a result of local instabilities [11]. Alternatively, fractal domains can be observed between mutant sectors arising from genetic differences in the population [12], where mutants with a fitness advantage grow faster and gain greater access to nutrients at the periphery of the colony [13]. Metabolic interactions between isogenic strains can lead to different patterns of self-organisation, from uniform radial expansion to the formation of dendritic niches at the colony edge [14].

Fractal boundaries have also been observed in *E. coli* cocultures of cross-feeding strains, where the type of social interaction determines the level of spatial mixing between the two strains [15]. The degree of self-similarity across the boundaries, usually measured through the fractal dimension, depends on the properties of the constituent cells.

The network of nutrient-transporting channels in *E. coli* biofilms [16] also exhibits a complex 3D morphology. The spatial structure of these emergent channels bears a striking resemblance to the fractal boundary patterning described in multi-strain co-culture biofilms, although these channels are not occupied by cells. In previous work, we measured the width of individual channels at different locations within the biofilm and showed that the channel architecture is affected by environmental growth conditions [17], but a quantification of channel morphology at the whole-biofilm scale has not been performed to date.

In this study, we quantify the morphological complexity of *E. coli* nutrient-transporting channels using fractal analysis of confocal microscopy images of biofilms. We hypothesised that their fractal nature would be affected by the shape of the constituent cells, because channel formation is an emergent property of biofilm growth. To test this hypothesis, we selected three *E. coli* mutant strains with altered cell shape phenotypes for morphological analysis. We show that the internal patterns formed by nutrient-transporting channels in *E. coli* have a fractal complexity comparable to that of computer-generated fractal images, and that this complexity is affected by cell shape and growth medium. We also report the specific case of Δ*ompR*, a wide-cell mutant whose biofilm morphological complexity is particularly affected by growth medium composition.

## 2. Results

### 2.1. Cell shape affects biofilm morphology

The single knockout mutants Δ*amiA*::kan, Δ*ompR*::kan and Δ*ydgD*::kan of the *E. coli* strain BW25113, hereby referred to as Δ*amiA*, Δ*ompR* and Δ*ydgD*, were chosen from the Keio collection for their altered cell shape phenotype (long, wide and wide respectively). The length and width of individual segmented cells were calculated for each strain from phase-contrast microscopy images (Supplementary Figure 1) and compared to the measurements of the isogenic parental strain BW25113 for quantitative phenotypic analysis. Δ*amiA* cells were on average 34% longer than the parental strain, whereas Δ*ompR* and Δ*ydgD* cells were on average 48% and 23% wider than the parental strain respectively.

After single-cell phenotypic characterisation, the three mutant strains were grown into mature biofilms on both LB (rich) and M9/glucose (minimal) solid growth medium (Figure 1). Confocal microscopy of these biofilms revealed a complex network of intra-colony channels, as previously reported for *E. coli* JM105 [16], [17]. Biofilms formed on minimal medium exhibited sectoring, with large areas of low fluorescence intensity across the biofilm volume, and channels appeared less morphologically complex than in their rich medium counterparts. This was particularly evident in the Δ*ompR* strain, where channels expand radially outwards in approximately straight lines, reminiscent of strain boundaries observed between isogenic domains in biofilms formed by spherical mutants of *E. coli* [11]. On rich medium, channels had more complex arrangements - in particular, channels formed by the Δ*amiA* strain were made up of long (≥ 10 μm) segments, consistent with the long cell phenotype of the constituent cells. Channels formed by the Δ*ompR* mutant strain, on the other hand, appeared fragmented, likely owing to the constituent cells’ irregular morphology.

**Figure 1:**
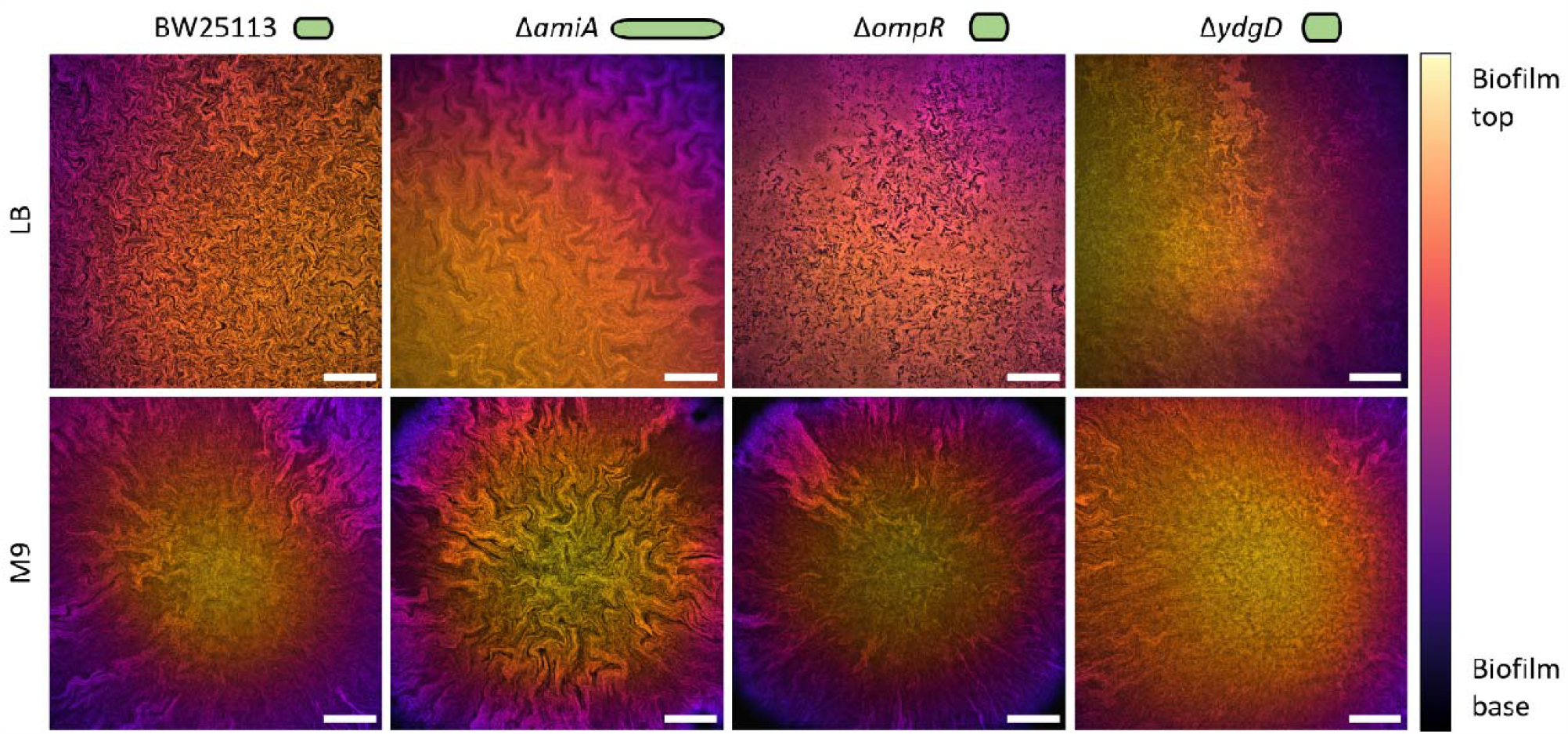
Representative confocal microscopy images showing the morphology of biofilms formed by the strains BW25113, Δ*amiA*, Δ*ompR* and Δ*ydgD* when growing on LB (rich) or M9 (minimal) solid medium. Images are projections of z-stacks acquired with 5 μm slice spacing, colour-coded by depth. The cell phenotype of each strain is shown as a green cartoon for reference. Scale bars: 200 μm.

Furthermore, despite both Δ*ompR* and Δ*ydgD* strains having a wide cell phenotype, their biofilms had different internal morphologies, especially on rich medium. Here Δ*ompR* formed darker channels, which were clearly distinguishable from the surrounding cells within the biofilm. On minimal medium, the central region of Δ*ydgD* biofilms appeared devoid of channels, whereas the edges showed radially expanding channels highly similar to those observed at the edges of Δ*ompR* biofilms. Enlarged regions of interest showing channel morphology are shown in Supplementary Figure 2.

### 2.2. Internal channel patterns in *E. coli* biofilms are fractal

The fractal complexity of internal biofilm patterns was quantified using relative differential box-counting (RDBC) dimension, which was calculated using the open-source plugin ComsystanJ on FIJI. Firstly, the plugin was used on four sets of computer-generated images with increasing spatial complexity (as described in the Methods), to obtain a correlation between visual complexity and RDBC dimension (Figure 2). Constant images had a RDBC dimension of 0 due to the lack of spatial features, whereas the RDBC dimension of images comprised of straight lines ranged between 2.114 ± 0.032 and 2.213 ± 0.043 (mean ± standard deviation). The RDBC dimension for computer-generated fractal images were calculated as between 2.329 ± 0.041 and 2.702 ± 0.047, and images of randomized features had a RDBC dimension equal to 2.959, close to the theoretically possible maximum of 3.000.

**Figure 2:**
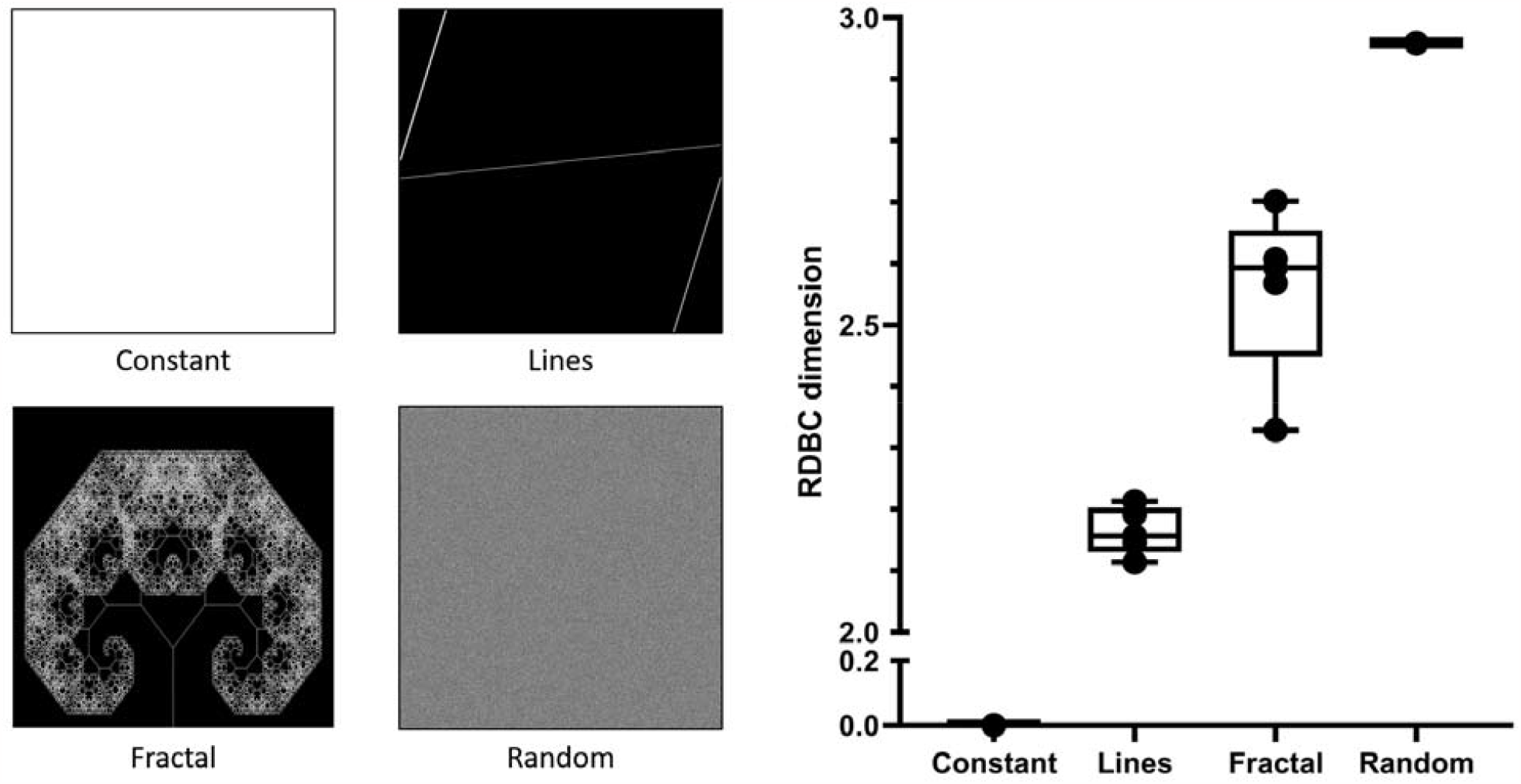
Representative images from the sets used for benchmarking of RDBC calculations (right panel). Each set consists of five images, whose fractal complexity is calculated using ComsystanJ. The constant, lines, and random images were generated in ComsystanJ, whereas the fractal images were obtained from open-source image repositories. Increasing complexity in the image sets is reflected by an increase in relative differential box-counting dimension, which ranges between 2.329 ± 0.041 and 2.702 ± 0.047 for fractal images.

After obtaining benchmark values for fractal complexity, confocal microscopy images of biofilms were analysed using the same image analysis pipeline to investigate the effect of cell shape and growth medium on the complexity of internal biofilm patterns (Figure 3). RDBC dimension values obtained from biofilm images fully aligned with those calculated for the set of computer-generated fractal images, with a minimum of 2.488 and a maximum of 2.620. This confirmed that channel structures in *E. coli* biofilms could be described using fractal geometry.

**Figure 3:**
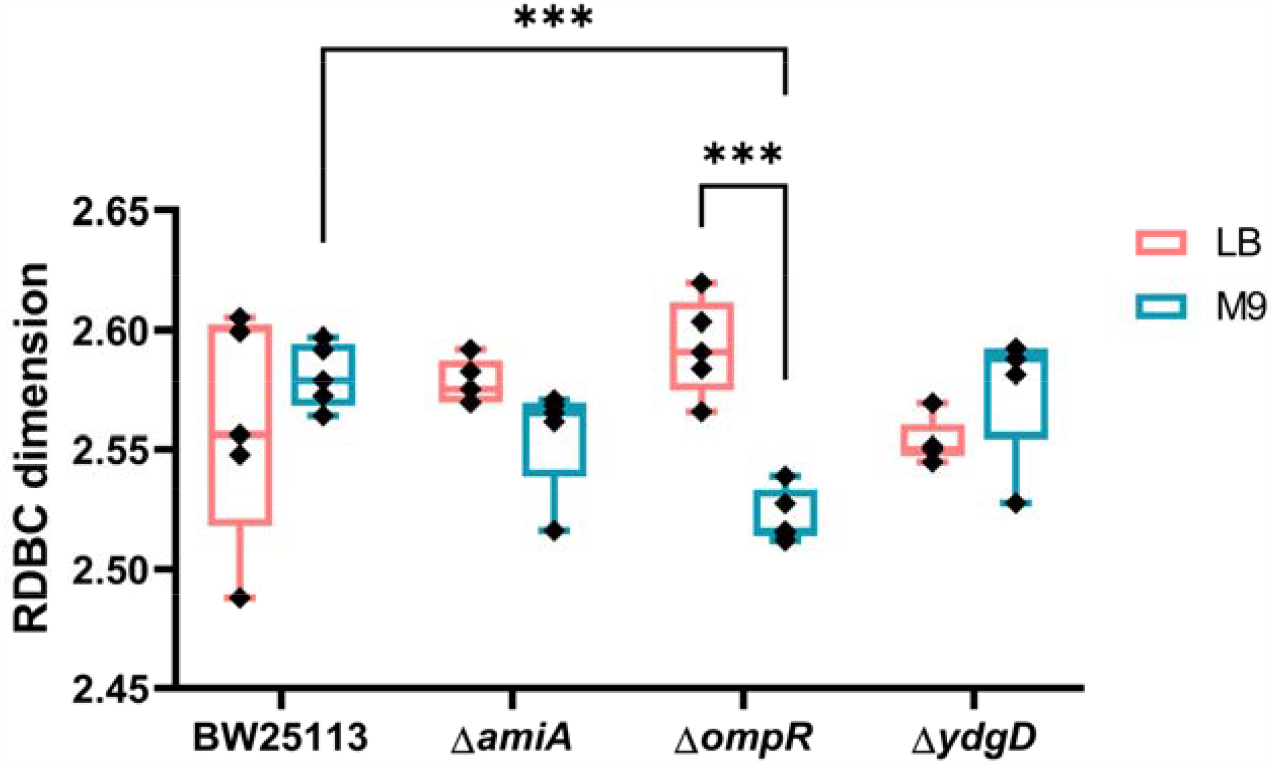
Box-counting dimension calculated from confocal micrographs of biofilms (n = 5 for each condition). On minimal medium (M9), biofilms formed by the Δ*ompR* mutant strain are less morphologically complex than those formed by the parental strain (p = 0.0005). For the Δ*ompR* mutant strain, the fractal complexity is also strongly affected by the growth substrate composition, with biofilms grown on rich medium (LB) showing significantly more complexity than those grown on minimal medium (M9, p = 0.0008). Average values are compared using one-way ANOVA tests.

On rich medium, biofilms formed by the Δ*amiA* mutant strain showed increased fractal complexity (RDBC = 2.578 ± 0.009) when compared to biofilms formed by Δ*ydgD* (RDBC = 2.553 ± 0.009, p = 0.0277). This reflects the peculiar channel architecture exhibited by Δ*amiA* biofilms grown on LB medium (Figure 1). On minimal medium, the parental strain BW25113 formed more complex biofilms (RDBC = 2.581 ± 0.014) than the Δ*ompR* mutant (RDBC = 2.522 ± 0.011, p = 0.0005). Biofilms formed by the mutant Δ*ydgD* were also less complex (RDBC = 2.576 ± 0.028) than those formed by Δ*ompR* (p = 0.0013), despite both strains having the same wide cell shape phenotype.

As shown in the image data presented in Figure 1, growth on minimal media was objectively correlated with large, dark sectors within the biofilm, which we hypothesised would result in a lower fractal complexity. However, after quantifying channel morphology through RDBC dimension and comparing each strain grown on the two different growth media we observed no statistical difference. An important exception to this observation was the Δ*ompR* mutant strain, for which growth in minimal medium was associated with significantly lower morphological complexity than growth in rich medium (p = 0.0008). This finding is investigated in more detail in the following sections.

### 2.3. Growth medium composition affects the complexity of Δ*ompR* biofilms

We further investigated the role of medium composition on Δ*ompR* biofilm internal morphology by comparing biofilms grown on LB and M9 substrates with different chemical compositions (Figure 4). The RDBC dimension of biofilms grown in rich medium decreased considerably when the substrate was made softer by halving the amount of agar (p = 0.0117). On minimal medium, on the other hand, growth on soft (1% agar) substrates was associated with a significant increase in RDBC dimension, comparable to that of biofilms grown on rich medium (p = 2.39 ⍰ 10^−7^).

**Figure 4:**
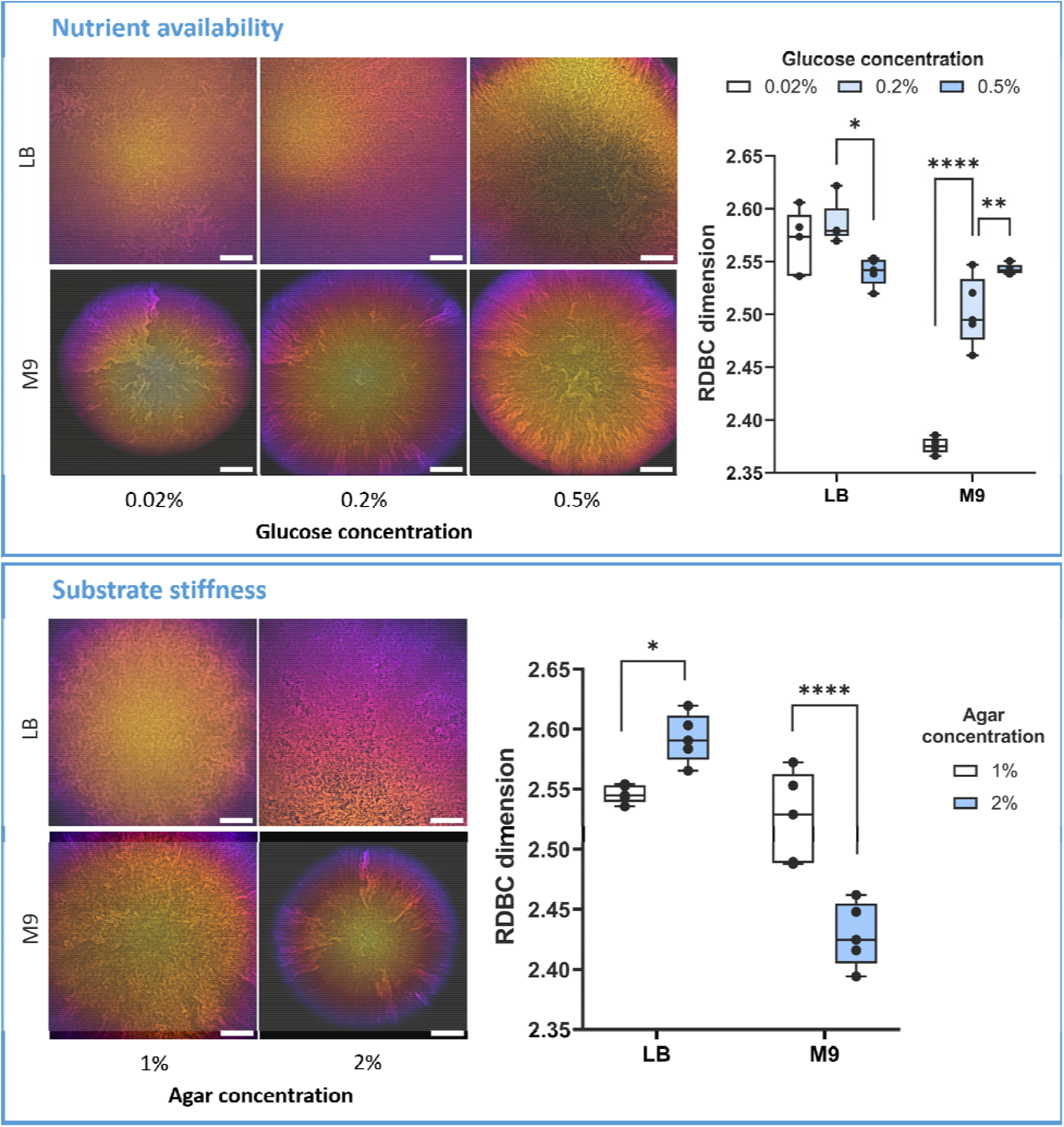
Morphology and fractal complexity of biofilms formed by the Δ*ompR* mutant strain grown on solid substrates with different glucose and agar concentrations. Glucose concentration in the medium is particularly important for the complexity of biofilms grown on minimal media (M9): RDBC dimension increases when glucose levels are increased from 0.02% to 0.2% (p = 4.81 ⍰ 10^−11^) and from 0.2% to 0.5% (p = 0.0478). On rich medium (LB), conversely, the addition of glucose does not significantly affect biofilm fractal complexity: only a small reduction in RDBC dimension is observed between 0.2% and 0.5% added glucose (p = 0.0166). Finally, a reduction in substrate stiffness from 2% to 1% agar concentration leads to a decrease in complexity on rich medium (p = 0.0365) and an increase in complexity on minimal medium (p = 2.39 ⍰ 10^−7^). Average values are compared using one-way ANOVA tests. Scale bars: 200 μm.

Furthermore, increasing glucose amounts in minimal medium led to an increase in RDBC from 2.376 ± 0.007 (0.02% glucose) to 2.503 ± 0.032 (0.2% glucose, p = 4.81 ⍰ 10^−11^) to 2.543 ± 0.005 (0.5% glucose, p = 0.0478). In Δ*ompR*, this increase in RDBC dimension coincided with the gradual disappearance of colony sectoring, brought by the increase in nutrient levels. As expected, the addition of the same amounts of glucose to rich medium did not lead to any significant changes in biofilm morphological complexity, with RDBC dimensions (2.520 < RDBC < 2.622) comparable to those found in nominal rich medium with no glucose (2.566 < RDBC dimension < 2.620).

While the diameter of Δ*ompR* biofilms increased with glucose concentration and decreased with agar concentration, the presence of dark background in images of smaller biofilms did not significantly affect the resulting RDBC dimension. This was checked by digitally zooming into images of the biofilms grown on 0.02% glucose (which had the smallest base area) until they filled the field of view of the images, and by successively calculating their RDBC dimension. The resulting RDBC value was calculated as 2.373 ± 0.006, a decrease of only 0.1% compared to the original biofilm images (Supplementary Figure 3).

### 2.4. Osmotic stress partially reduces biofilm complexity in Δ*ompR*

Because of the role of *ompR* in osmotic stress response, the fractal complexity of biofilms formed by the mutant strain Δ*ompR* was studied on LB and M9/glucose medium with and without salt (Figure 5). A reduction in RDBC dimension was observed when the osmolality of the medium was reduced by removing salt, both in rich medium (p = 4.88 ⍰ 10^−5^) and in minimal medium (p = 0.0209). This suggested that osmotic stress might control the morphology of biofilms formed by the Δ*ompR* mutant strain.

**Figure 5:**
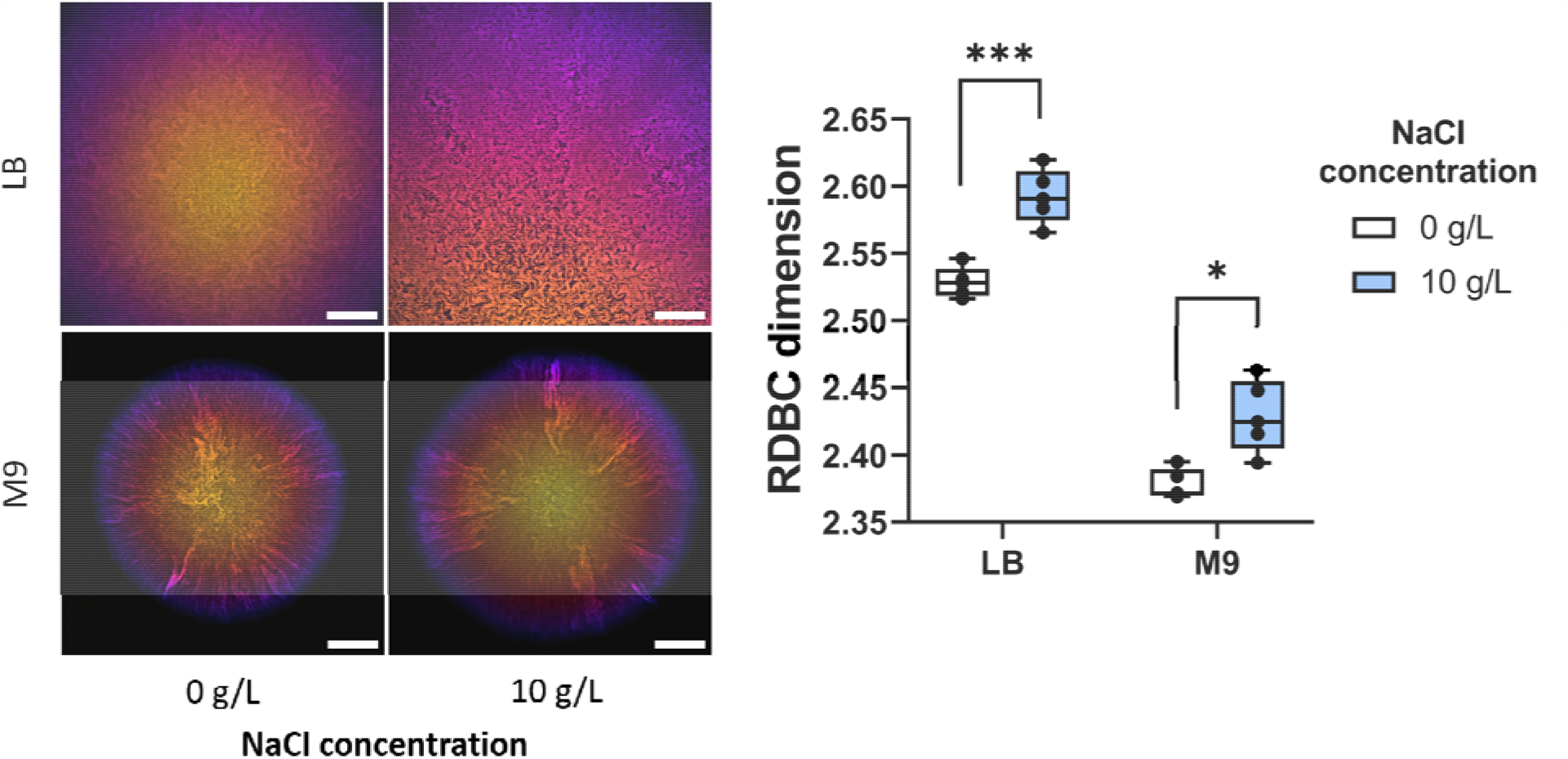
Morphology and fractal complexity of biofilms formed by the Δ*ompR* mutant strain grown on LB and M9/glucose solid substrates with and without NaCl. The presence of 10 g/L NaCl is associated with an increase in biofilm morphological complexity for both rich and minimal media (p = 7.34 ⍰ 10^−4^ and p = 0.0209 respectively), as indicated by the RDBC dimension metric.

For this reason, the role of osmotic stress on the growth of the Δ*ompR* mutant strain was investigated at both the cellular level and at the biofilm level. This was achieved by measuring changes in cell length and width and by measuring the RDBC dimension of biofilm images grown in solid media with different osmolalities, respectively. Changes in growth medium osmolality were induced by adding different amounts of iodixanol, a compound originally designed for density gradient preparations that cannot be metabolised by the bacteria [18]. Iodixanol toxicity in both the parental strain BW25113 and the Δ*ompR* mutant was checked by growth curve experiments (Supplementary Figure 4). At the cellular level, an increase in osmotic stress in the liquid growth media led to a reduction in cell size for both strains (Supplementary Figure 5). A significant reduction in cell length with increasing iodixanol concentration was observed in both strains: from 3.520 ± 0.827 μm to 2.205 ± 0.516 μm for BW25113 (p = 1.32 ⍰ 10^−167^), and from 3.878 ± 1.133 μm to 2.685 ± 0.657 μm for Δ*ompR* (p = 6.74 ⍰10^−40^). By contrast, cell width reduction with increasing iodixanol concentration was marginal for BW25113 (from 1.001 ± 0.099 μm to 0.970 ± 0.098 μm, p = 8.57 ⍰ 10^−8^), whereas it was almost five times higher for the Δ*ompR* strain (from 1.335 ± 0.219 μm to 1.147 ± 0.149 μm, p = 1.02 ⍰ 10^−13^).

In biofilms, an increase in iodixanol concentration in the solid growth substrate was associated with a small overall reduction in RDBC dimension for both strains (Figure 6). This loss of internal complexity could occur because planktonic growth is limited in high osmolality medium (Supplementary Figure 4), or it could be a result of the cell size reduction following osmotic stress (Supplementary Figure 5). Alternatively, it could be a pleiotropic effect caused by the presence of iodixanol in the growth medium.

**Figure 6:**
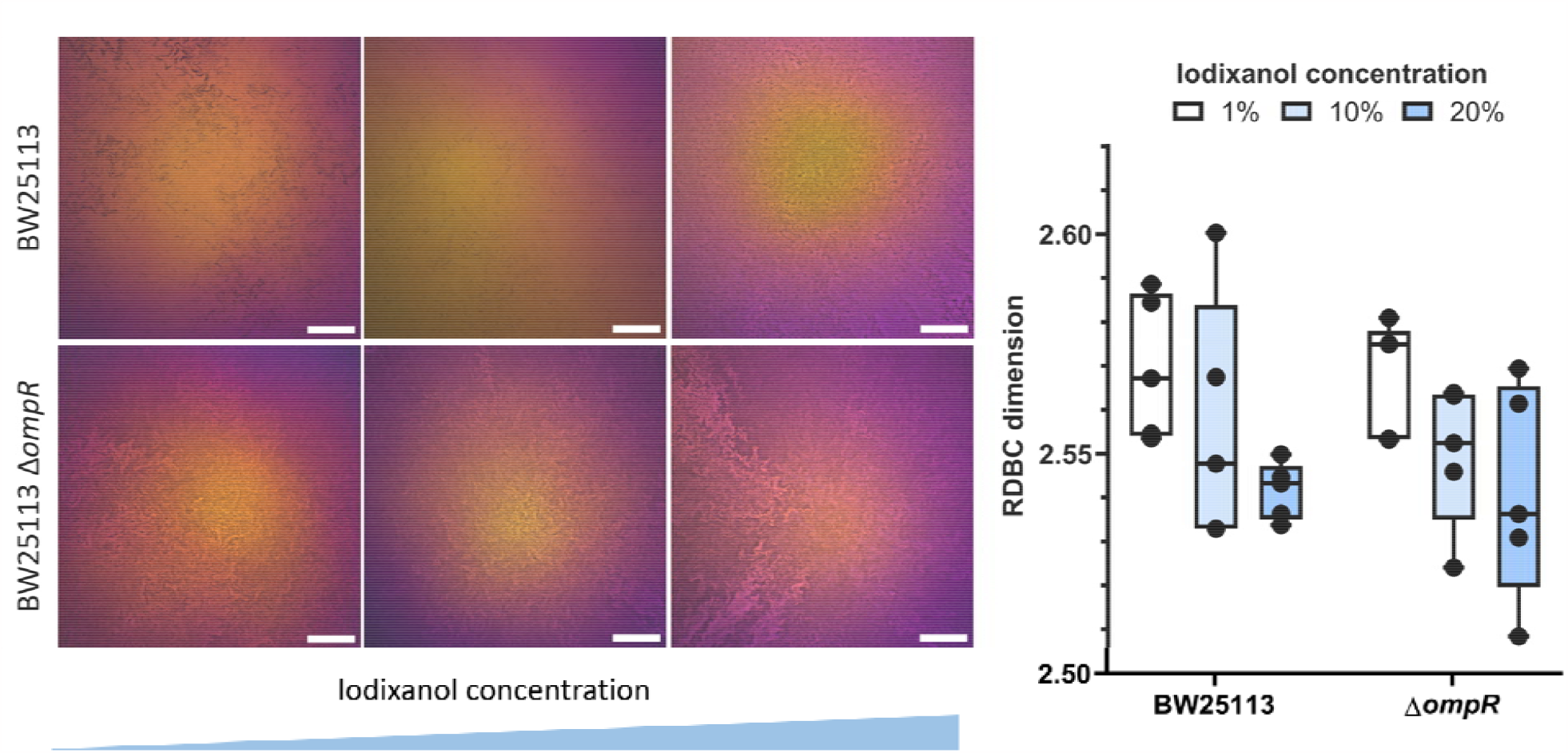
Comparison of fractal morphology between biofilms formed by the parental strain BW25113 and the mutant strain Δ*ompR* on solid substrates with increasing iodixanol concentrations (1%, 10% and 20% v/v). Biofilm images are colour-coded by depth. Average RDBC dimension decreases with increasing amounts of iodixanol and is similar between the two strains for each iodixanol concentration. Scale bars: 200 μm.

## 3. Discussion

The formation of intra-colony channel networks is an emergent property of *E. coli* biofilms, where these channels constitute a novel nutrient uptake system [16]. The chemical makeup of the growth environment is known to affect the density and the size of these channels within the biofilm [17], however the factors controlling channel morphogenesis are currently unknown. A method for the systematic, quantitative characterisation of channel morphology is therefore needed to elucidate the mechanisms governing channel formation.

In this study, we analysed the effect of cell shape mutations and growth medium composition on the morphological complexity of *E. coli* biofilms using confocal microscopy and image analysis. We showed that the fractal morphology of *E. coli* nutrient transport channels was comparable to that of computer-generated fractals for all cell shape mutants and growth conditions. While cell shape affected the overall biofilm morphology, this did not lead to a significant change in fractal complexity, as described by the RDBC dimension metric. However, the wide-cell mutant strain Δ*ompR* formed biofilms with a higher RDBC dimension on nutrient excess than on nutrient limited conditions.

To investigate the cause of this phenomenon, we examined the effect of growth medium composition on Δ*ompR* biofilm fractal complexity. We found that glucose concentration was particularly important in minimal medium, where it constituted the sole carbon source. Increasing glucose concentrations in minimal medium led to proportionally more complex biofilm channel organisation, eventually reaching a RDBC dimension value close to that of biofilms grown on rich media. A similar effect was previously observed in *E. coli* JM105 mini-Tn7-*gfp* biofilms, which developed more complex channel patterns when grown on minimal medium with an excess of glucose compared to glucose-limited substrates [17]. Furthermore, we found that growth on rich soft substrates reduced biofilm morphological complexity, whereas growth on soft minimal substrates increased it. This is in contrast with previous findings on *E. coli* JM105 mini-Tn7-*gfp*, where channels from biofilms grown on rich, soft substrates were densely packed [17].

We investigated the morphology of Δ*ompR* cells and biofilms under growth conditions with varying osmolality owing to the role of OmpR in regulating the osmotic stress response in *E. coli* [19]. Gram-negative bacteria such as *E. coli* respond to an increase in external osmotic pressure by accumulating solutes inside the cell, and by pumping out water through efflux [20]. Hyperosmotic shock also leads to a sudden cell volume shrinkage [21], followed by a gradual recovery [22], and is associated with a reduction in cell elongation rate [23]. We investigated the effect of medium osmolality on biofilm morphology by comparing RDBC dimension values between biofilms grown on solid substrates with and without NaCl, and we found that low osmolality was associated with a lower fractal complexity on both rich and minimal media. Because NaCl affects not only medium osmolality, but also nutrient metabolism [24], we then used iodixanol for subsequent osmotic stress experiments. The observed reduction in Δ*ompR* cell width in high osmolality medium indicates a possible partial reversion to the parental phenotype, which could equally be driven by *ompR* suppressor mutations. This phenotypic reversion in cell shape may also explain the similarity in absolute RDBC dimension values between BW25113 and Δ*ompR* biofilms at each iodixanol concentration.

The extraction of fractal geometry parameters from biofilm micrographs (see for example [25]) is usually achieved by image thresholding, a type of image segmentation which isolates an object from its background depending on its grayscale value [26]. While this method can accurately identify biofilm outlines, it is not effective for the detection of internal channel networks, where difference in grayscale values between the channels and the rest of the biofilm is small. In our work, this obstacle is overcome using the fractal analysis software ComsystanJ. The use of fractal geometry has proven to be a simple yet powerful method for the quantification of *E. coli* biofilm internal channels formed by cell shape mutants on different growth substrates. Our analysis could help identify the factors controlling the formation and development of nutrient-transporting channels in *E. coli*, and it could be readily applied to other microbial species exhibiting complex biofilm internal patterns.

## 4. Materials and methods

### 4.1. Strains and media

The bacterial strains used in this work (Supplementary Table 1) were obtained from the Keio collection [27], a single-gene knockout library of all nonessential genes in the *E. coli* K-12 strain, BW25113 [28]. The mutants Δ*amiA*::kan, Δ*ompR*::kan and Δ*ydgD*::kan of the *E. coli* strain BW25113, hereby referred to as Δ*amiA*, Δ*ompR* and Δ*ydgD*, were selected for their modified cell phenotype, and their single-gene deletions were verified by PCR and DNA amplicon sequencing (Supplementary Table 2). The strains were transformed by electroporation with the plasmid pAJR145, which is derived from the plasmid pACYC184 and encodes the constitutively-expressed green fluorescent protein (GFP) transcriptional fusion *rpsM::gfp+* [29]. Prior to transformation, liquid cultures of each strain were made electrocompetent through three ice-cold 10% glycerol washes.

Liquid cultures were grown overnight in a 37°C aerated incubator while shaking at 250 rpm, in Miller Lysogeny Broth (LB) [30] with the addition of 25 μg/mL chloramphenicol to maintain GFP fluorescence from the pAJR145 plasmid. Keio mutant strains were also grown with the addition of 50 μg/mL kanamycin. M9 minimal medium salts [30] were prepared as a 5× solution, then diluted to 1× with distilled deionised water and supplemented with 1 mM MgSO_4_·7H_2_O, 0.2% (w/v) glucose and 0.00005% (w/v) thiamine. Solid substrates were prepared by adding 20 g/L of agar and were grown in a 37°C static aerated incubator.

Solid growth substrates were also prepared with different nutritional profiles and agar concentrations to quantify the resulting change in internal morphology of Δ*ompR* biofilms. No-salt agar substrates were prepared without NaCl, and soft agar substrates were prepared by reducing the amount of agar to 10 g/L. D-Glucose was added to both LB and M9 media to final concentrations of 0.02% (w/v), 0.2% (w/v) or 0.5% (w/v).

### 4.2. Growth characterisation of BW25113 and Δ*ompR* in high osmolality medium

The growth of the parental strain BW25113 and of the mutant strain Δ*ompR* in liquid medium were characterised using a plate reader. LB broth was prepared with iodixanol concentrations of 0%, 1%, 2%, 5%, 10% and 20% (v/v) by adding appropriate amounts of an OptiPrep 60% (w/v) iodixanol stock solution (Sigma-Aldrich, USA). Ten wells of a 96-well plate were then filled with 200 μL of medium with each iodixanol concentration (corresponding to five biological replicates for each strain). Overnight cultures of BW25113 and of the Δ*ompR* mutant strain were prepared in LB broth as described above and were diluted to a starting OD_600_ of 0.01. The plate was then loaded with the lid on onto a pre-warmed (37°C) Synergy HT plate reader (BioTek, USA), where the OD_600_ of the cultures was measured every 15 minutes for 24 hours while the plate was shaking continuously at medium speed.

Growth curves were plotted in Prism by exporting data from the Gen5 microplate software (BioTek, USA), with the y axis plotted on a logarithmic scale. Specific growth rates were calculated as the slope of the linear region of the semi-logarithmic plot using the equation

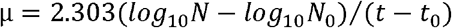

*N*_0_ and *N* are the cell numbers at the beginning and at the end of the exponential growth phase, respectively, and correspond to optical density readings, and *t*_0_ and *t* are the times at which exponential growth phase starts and ends, respectively [31].

To compare cell shape phenotype before and after growth in high osmolality medium, BW25113 and Δ*ompR* were first grown as liquid cultures overnight in LB broth. They were then diluted 1:100 in both LB broth and LB broth with 20% (v/v) iodixanol with LB salts increased proportionally, and were incubated further in a 37°C aerated incubator while shaking at 250 rpm until mid-exponential growth phase, before imaging using phase contrast microscopy as described below. Solid LB medium plates were prepared with iodixanol concentrations of 1%, 10% and 20% (v/v), and used as substrates for the growth of BW25113 and Δ*ompR* biofilms.

### 4.3. Single-cell phase contrast microscopy

For single-cell imaging, overnight cultures of each strain in LB broth were diluted 1:100 and incubated for two hours until they reached mid-exponential growth phase (OD_600_ = 0.4-0.6). Imaging slides were prepared by sandwiching 1 mL of molten 1% agarose between two microscope slides and letting it solidify at room temperature. After removing the top slide, 10 μL of liquid culture was spotted onto the solidified agarose pads, and a coverslip was added prior to imaging. Single-cell phase contrast images were acquired for both non-fluorescent strains (BW25113, BW25113 Δ*amiA*::kan, BW25113 Δ*ompR*::kan, BW25113 Δ*ydgD*::kan) and fluorescent strains (BW25113 / pAJR145, BW25113 Δ*amiA*::kan / pAJR145, BW25113 Δ*ompR*::kan / pAJR145, BW25113 Δ*ydgD*::kan / pAJR145), to ensure that the addition of chloramphenicol required for the maintenance of the pAJR145 plasmid did not affect the cell phenotype of each strain.

Single cell imaging was carried out using an Eclipse E600 upright widefield microscope (Nikon, Japan) in phase contrast mode equipped with a 100x/1.30 DLL oil immersion lens (Nikon, Japan). Illumination was provided by a halogen lamp and the image was detected using a Hamamatsu ORCA 100 digital camera (Hamamatsu, Japan).

### 4.4. Biofilm confocal microscopy

The GFP-expressing strains BW25113 / pAJR145, BW25113 Δ*amiA*::kan / pAJR145, BW25113 Δ*ompR*::kan / pAJR145 and BW25113 Δ*ydgD*::kan / pAJR145 were grown into mature biofilms on agar substrates in a sterile 3D-printed plastic specimen holder, as described previously [16]. Biofilms were grown for 24 hours when using LB medium, and for 48 hours when using M9 medium.

Mature biofilm images were acquired on an Olympus IX81 microscope coupled to a FluoView FV1000 confocal laser scanning unit (Olympus, Japan). Fluorescence from GFP was excited using a 488 nm argon laser (GLG3135, Showa Optronics, Japan) and was detected by a PMT with a spectral detection window set between wavelengths of 510 and 560 nm. Samples were imaged using a 10x/0.4 N.A. air objective lens for resolving intra-colony channels.

Three-dimensional z-stacks of biofilms grown on LB and M9 media were acquired with a slice spacing of 5 μm for Nyquist sampling in the axial dimension. Five different colony biofilms were imaged for each condition.

### 4.5. Image analysis

Phase-contrast images of both fluorescent and non-fluorescent cells were first pre-processed in FIJI using the background subtraction tool with a rolling ball radius of 15 pixels, with the “light background” option selected. The images were then analysed using the FIJI plugin MicrobeJ [32] to obtain cell length and width measurements. Segmentation was performed using MicrobeJ with the default settings for a bright background and the “medial axis” mode of detection. The following changes to the default parameter ranges were made: area [1-max] μm^2^; width [0-max] μm with variation [0-0.2]; sinuosity [1-1.2]; angularity [0-0.5] rad; solidity [0.9-max]. The default “advanced” parameters were changed to have an area cut-off of 1000, and a count cut-off of 250. The options “exclude on edges” and “shape descriptors” were selected. Phase contrast images were analysed (n = 10 for each strain), with a total number of analysed cells between 202 and 665 (non-fluorescent cells) and between 90 and 524 (fluorescent cells).

Biofilm image z-stacks were displayed as hyperstacks, and colour-coded by depth using the “fire” lookup table on FIJI. Biofilms images were contrast-adjusted where needed for presentation purposes, using Contrast Limited Adaptive Histogram Equalization (CLAHE, [33]) available in FIJI with block size 60, maximum slope 3 and 256 histogram bins. Regions of interest (ROIs) in biofilm images were despeckled using the “noise – despeckle” function in FIJI.

### 4.6. Fractal complexity quantification

Fractal complexity was quantified using the FIJI plugin ComsystanJ (Complex Systems Analysis for ImageJ) [34], version 1.0.0 (https://github.com/comsystan/comsystanj). The “2D image” version of the plugin was used on the images, all of which had a size of 2048 × 2048 pixels. Box-counting fractal dimension was calculated using the Relative Differential Box Counting (RDBC) algorithm developed by Jin et al. [35] which uses a raster box scanning method. The plugin was run with 12 boxes and 1-12 regressions.

Firstly, four sets of sample images were analysed. Three of the sets were generated as grayscale images through ComsystanJ using the “2D image Image Generator” option, with image types “Constant”, “Fractal random shapes – Lines” and “Random”. Constant images had a constant pixel intensity value (256) throughout each image. Line images were made of light (pixel intensity = 256) lines with varying thickness randomly intersecting on a dark (pixel intensity = 0) background. Fractal images were obtained from the “Wikimedia commons” open-source image repository [36]–[40]. Finally, in Random images every pixel had a random intensity value (between 0 and 256).

For biofilm fractal pattern analysis, z-stacks were converted to 8-bit. No contrast-adjustment was used on the images prior to analysis.

### 4.7. Statistical analysis

Statistical tests were carried out on Prism version 8.0.2 (GraphPad Software, USA). Cell measurements (length and width) of the mutant strains were compared to those of the parental strain BW25113 using a Kruskal-Wallis multiple comparisons test. One-way ANOVA tests were used to compare box-counting dimensions of biofilms formed by different strains in the same media and also to compare box-counting dimensions of the same strain grown in different media.

In the main text and figure captions, values are presented as mean ± standard deviation. In all plots, produced using Prism version 8.0.2, p-values are presented as * (p < 0.05); ** (p < 0.005); *** (p < 0.0005); **** (p < 0.0001), with specific p-values written in the figure captions.

## 5. CRediT authorship contribution statement

**Beatrice Bottura**: Conceptualisation, Methodology, Validation, Formal Analysis, Investigation, Resources, Data Curation, Writing – Original Draft, Writing – Review & Editing, Visualisation.

**Liam M. Rooney**: Conceptualisation, Methodology, Resources, Writing – Review & Editing.

**Morgan Feeney**: Methodology, Supervision, Writing – Review & Editing

**Paul A. Hoskisson**: Conceptualisation, Methodology, Writing – Review & Editing, Resources, Supervision, Project Administration, Funding Acquisition.

**Gail McConnell**: Conceptualisation, Methodology, Writing – Review & Editing, Resources, Supervision, Project Administration, Resources, Funding Acquisition.

## 7. Acknowledgements

We thank Dr Manuel Banzhaf (University of Birmingham) for the gift of the Keio collection mutants. We also thank Dr Ainsley Beaton (John Innes Centre, Norwich), Dr Rebecca McHugh and Dr David Mark (University of Glasgow), Robyn Braes, Elmira Mohit and James Croxford (University of Strathclyde) for helpful discussions.

Beatrice Bottura is supported by a University of Strathclyde Student Excellence Award. Liam Rooney is supported by The Leverhulme Trust. Gail McConnell is funded by the Medical Research Council (MR/K015583/1), the Biological Sciences Research Council (BB/T011602/1 and The Leverhulme Trust. Paul A. Hoskisson is supported by BBSRC (BB/T001038/1 and BB/T004126/1), MRC (MR/V011499/1), The Leverhulme Trust and the Royal Academy of Engineering Research Chair Scheme for long term personal research support (RCSRF2021\11\15).

For the purpose of open access, the author(s) has applied a Creative Commons Attribution (CC BY) licence to any Author Accepted Manuscript version arising from this submission.

