## Supplementary Information for "Fractal complexity of Escherichia coli nutrient transport channels is influenced by cell shape and growth environment"

### 1. Supplementary information

#### 1.1. Bacterial strains

| **Strain** | **Genotype** | **Reference** |
| --- | --- | --- |
| BW25113 | F-, *Δ(araD-araB)567*, *ΔlacZ4787(*::rrnB-3*), λ-, rph-1, Δ(rhaD-rhaB)568, hsdR514* | [1] |
| BW25113 / pAJR145 |  | This study |
| BW25113 Δ*amiA*::kan | BW25113 Δ*amiA764::kan* | [2] |
| BW25113 Δ*ompR*::kan | BW25113 Δ*ompR739::kan* | [2] |
| BW25113 Δ*ydgD*::kan | BW25113 Δ*ydgD746::kan* | [2] |
| BW25113 Δ*amiA*::kan / pAJR145 |  | This study |
| BW25113 Δ*ompR*::kan / pAJR145 |  | This study |
| BW25113 Δ*ydgD*::kan / pAJR145 |  | This study |

Supplementary Table 1: Bacterial strains used in this work.

#### 1.2. Genetic verification of Keio mutants

The genotype of the Keio mutant strains Δ*amiA*::kan, Δ*ompR*::kan and Δ*ydgD*::kan was checked by PCR and DNA sequencing. Genomic DNA from each strain was purified using a Wizard Genomic DNA Purification Kit (Promega, UK) and was used as template in PCRs carried out with PCRBIO Taq Mix Red (PCR Biosystems, USA) following manufacturer’s instructions. The presence of the kanamycin resistance cassette was verified by using the k1 and k2 primers from [1], whereas the deletion of each gene was confirmed with gene-specific primers designed up- and down-stream of the genes *amiA*, *ompR* and *ydgD* (Supplementary Table 2).

| **Primer name** | **Primer sequence (5’ - 3’)** | **Tm (°C)** |
| --- | --- | --- |
| amiA_F | TCTCAACAGCAAACCGTCGT | 67 |
| amiA_R | GTTTAACCTGGTGTGCGTCG | 67 |
| ompR_F | TAGCTGGTGACGAACGTGAG | 67 |
| ompR_R | GCGAACAGCAAGGTGACGAT | 68 |
| ydgD_F | ACTTTCATCCCGTCCCGTCT | 68 |
| ydgD_R | ATTGGCCTGGTCTTGCTGTT | 67 |

Supplementary Table 2: Primers used for PCR amplification of the regions containing the genes of interest.

The deletion of each gene of interest and its substitution with a kanamycin resistance cassette were also confirmed by DNA sequencing of the chromosomal regions containing the genes of interest (Eurofins, Germany). This was achieved using purified PCR products obtained from the reactions described above, and the gene-specific primers from Supplementary Table 2.

#### 1.3. Single-cell phenotypic characterisation of cell shape mutants

Cell shape mutations are often achieved by deleting genes involved in cell wall synthesis [3]. In this work, we selected knockout mutants of the genes *amiA*, *ompR* and *ydgD* for cell shape and biofilm morphological analysis.

Deletion of the *amiA* gene leads to incomplete septum cleavage in the peptidoglycan cell wall [4]. While it has been reported that 5-10% of Δ*amiA* mutant cells of *E. coli* grow as chains [5], this is only the case in stationary growth phase [6]. In our imaging experiments, performed at mid-exponential growth phase, Δ*amiA* cells have an average length of 4.614 ± 1.640 µm, which is longer than that of the parental strain, 3.441 ± 0.803 µm (p = 3.48 ⨯ 10^-11^).

Deletion of the *ompR* gene in *E. coli* inactivates the expression of the outer membrane porins OmpC and OmpF, which mediate the diffusion of small solutes across the outer membrane [7]–[9]. Δ*ompR* cells have above-average width of 1.582 ± 0.336 µm (compared to the parental strain’s 1.066 ± 0.112 µm, p = 5.16 ⨯ 10^-39^). Our increased cell width measured for the Δ*ompR* mutant contradicts with that calculated by French et al. in the same medium [10], who did not identify the deletion of *ompR* as a cause for cell phenotype change, but concurs with that calculated by Campos et al. [11]. This could be due to French’s use of a 2% glutaraldehyde fixatives (pH 6.8) prior to imaging. In fact, the addition of 2% glutaraldehyde is associated with up to two-fold increase in PBS osmolality [12], up to 2.5 times higher that of LB broth [13], and *E. coli* fixation in Chemicon is associated with an immediate decrease in cell width of approximately 15% [14]. Interestingly, the Δ*ompR* mutant exhibits an irregular, bulgy cell shape phenotype, similar to that observed by Ranjit and Young during growth of lysozyme-induced spheroplasts of *E. coli* MG1655 with the same Δ*ompR* mutation [15]. Nonetheless, this morphology was not observed in the two phenotypic studies mentioned above. We do not ascribe this to differences in immobilisation method (agarose pads for our study and Campos’; optically clear microplates for French’s), which is known not to affect cell morphology, changes in intercellular pH or growth in *E. coli* [16].

The gene *ydgD* is a putative periplasmic serine protease, and its deletion in *E. coli* is linked to increased sensitivity to some β-lactam antibiotics [17]. In our work, Δ*ydgD* cells have a wide phenotype, with average width 1.313 ± 0.221 µm (larger than the width of the parental strain, p = 3.81 ⨯ 10^-28^). This is consistent with reports in the Keio collection [2].

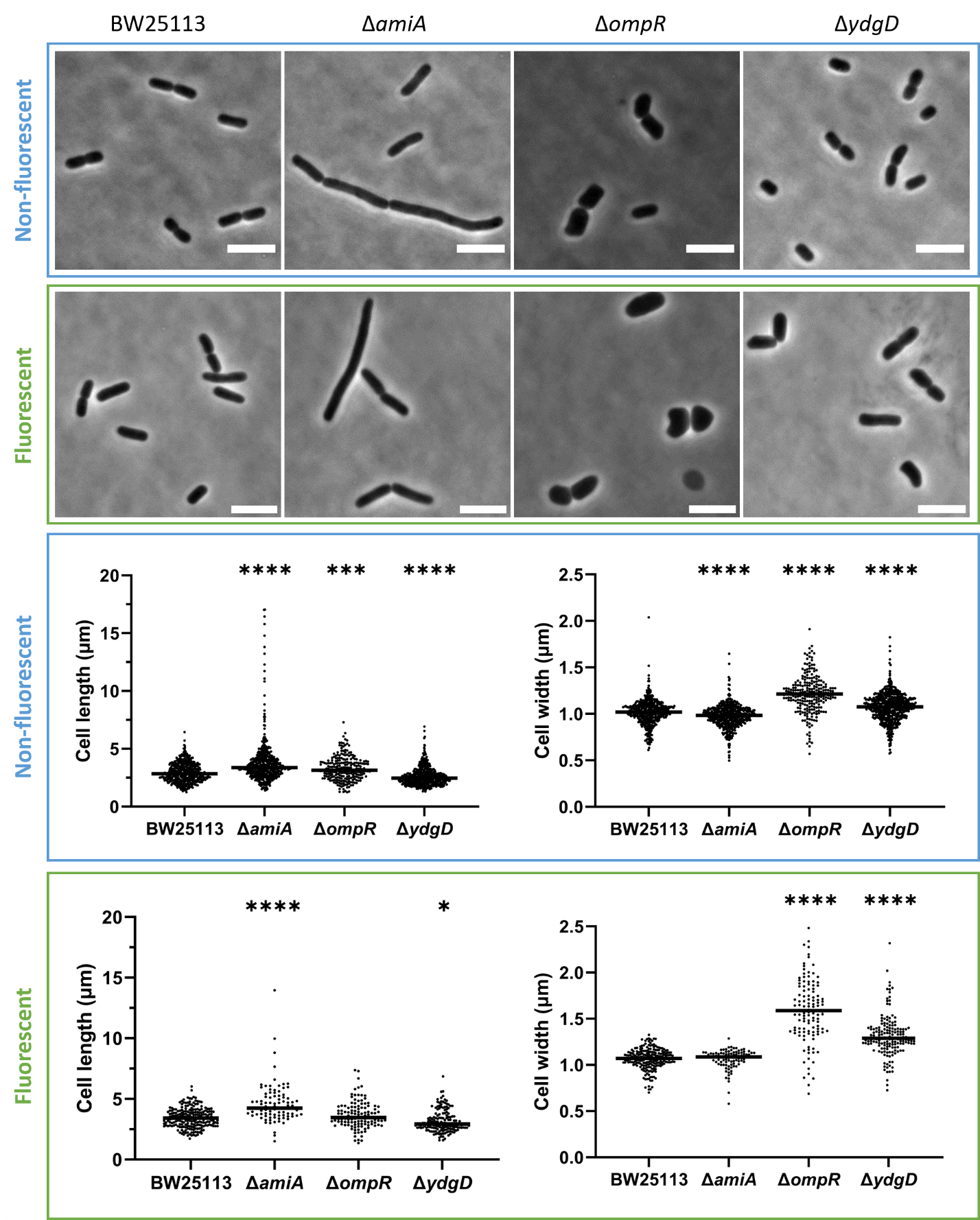

Supplementary Figure 1: (a) phase contrast single-cell images of the Keio parental strain BW25113 and of three cell-shape mutants, showing long (Δ*amiA*) and wide (Δ*ompR* and Δ*ydgD*) phenotypes. Images were acquired of both non-fluorescent and fluorescent strains. Scale bars: 5 µm.

(b) Cell length and width measurements for fluorescent (green panel) and non-fluorescent (blue panel) strains. Horizontal lines denote the median of each dataset, and asterisks represent statistical significance between each mutant strain and the parental strain, with average values compared with a Kruskal-Wallis statistical test. Δ*amiA* is long (p = 1.34 ⨯ 10^-18^, n = 562 non-fluorescent cells and p = 3.48 ⨯ 10^-11^, n = 90 fluorescent cells). Δ*ompR* is wide (p = 3.39 ⨯ 10^-42^, n = 238 non-fluorescent cells and p = 5.16 ⨯ 10^-39^, n = 160 fluorescent cells). Δ*ydgD* is short (p = 2.78 ⨯ 10^-11^, n = 618 non-fluorescent cells and p = 0.0134, n = 120 fluorescent cells) and wide (p = 1.84 ⨯ 10^-10^ for non-fluorescent cells and p = 3.81 ⨯ 10^-28^ for fluorescent cells).

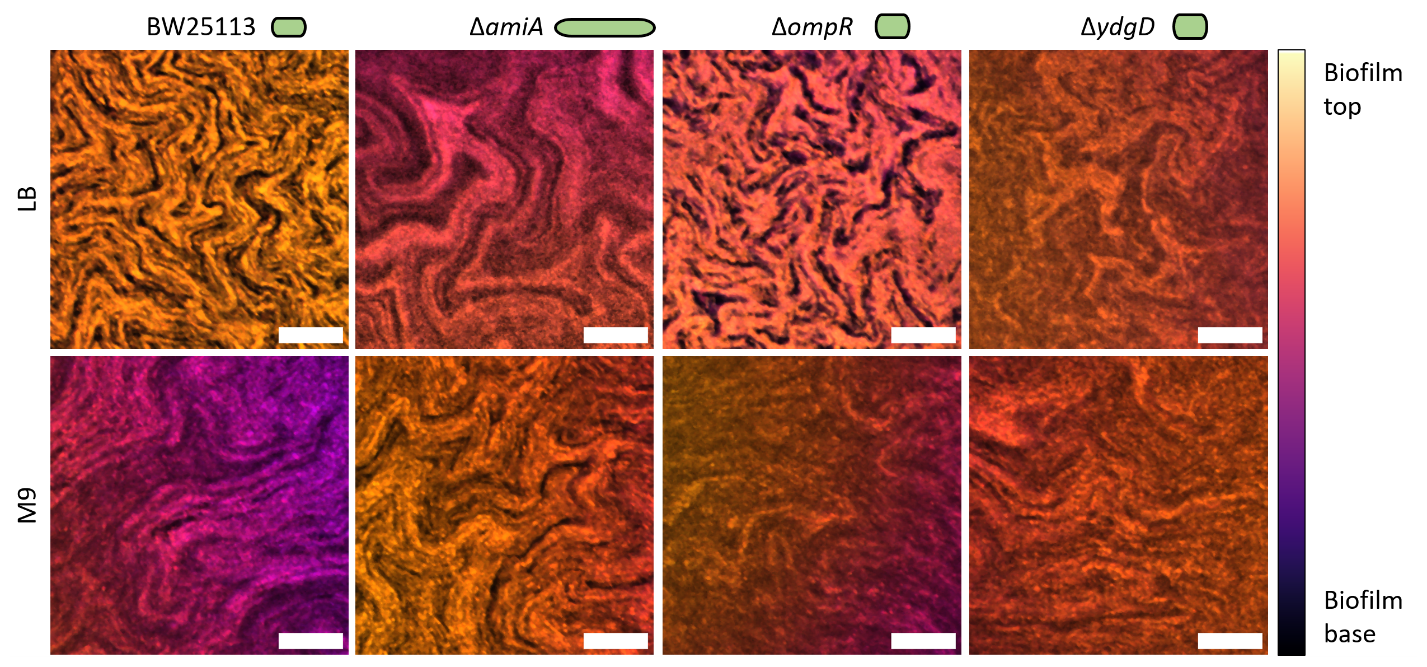

Supplementary Figure 2: ROIs of z-stack projections from Figure 1, showing the morphology of individual channels. Images are locally contrast-adjusted using CLAHE, and despeckled using FIJI for presentation purposes. The cell phenotype of each strain is shown as a green cartoon for reference. Scale bars: 50 µm.

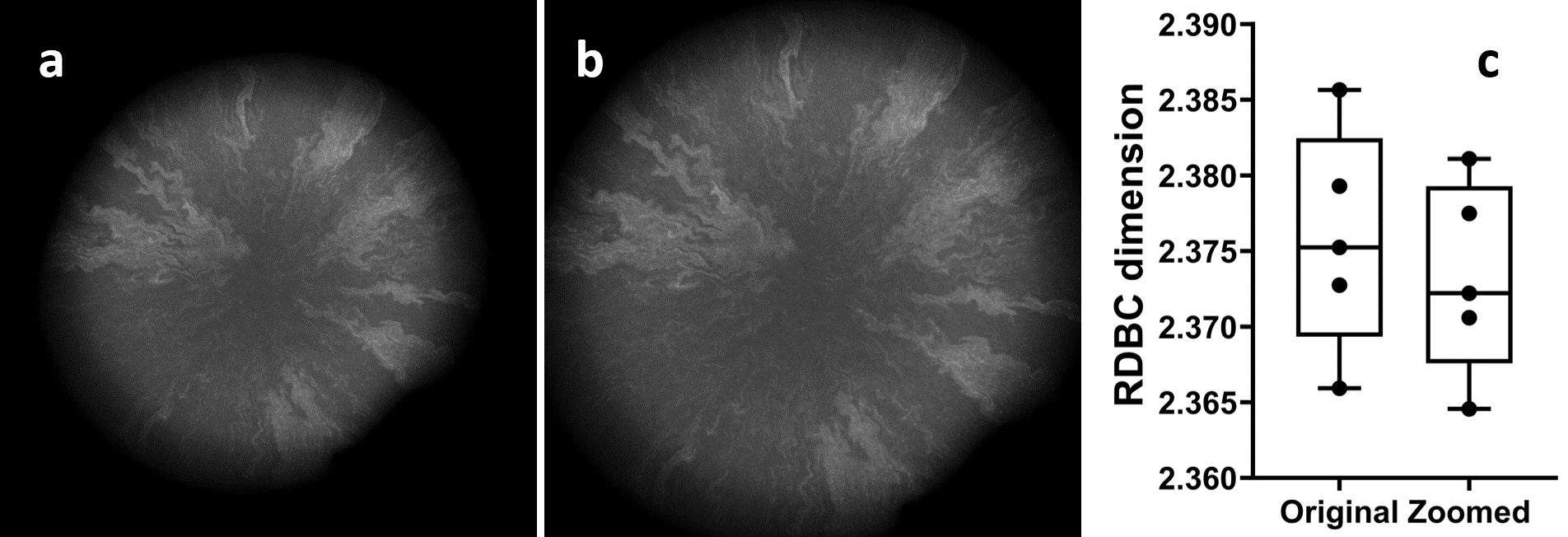

Supplementary Figure 3: Comparison of RDBC dimension values (c) calculated for original Δ*ompR* biofilm images (a) and the same images zoomed to fill the image space and re-scaled to 2048 × 2048 pixels (b). The resulting average decrease in RDBC is only 0.1%.

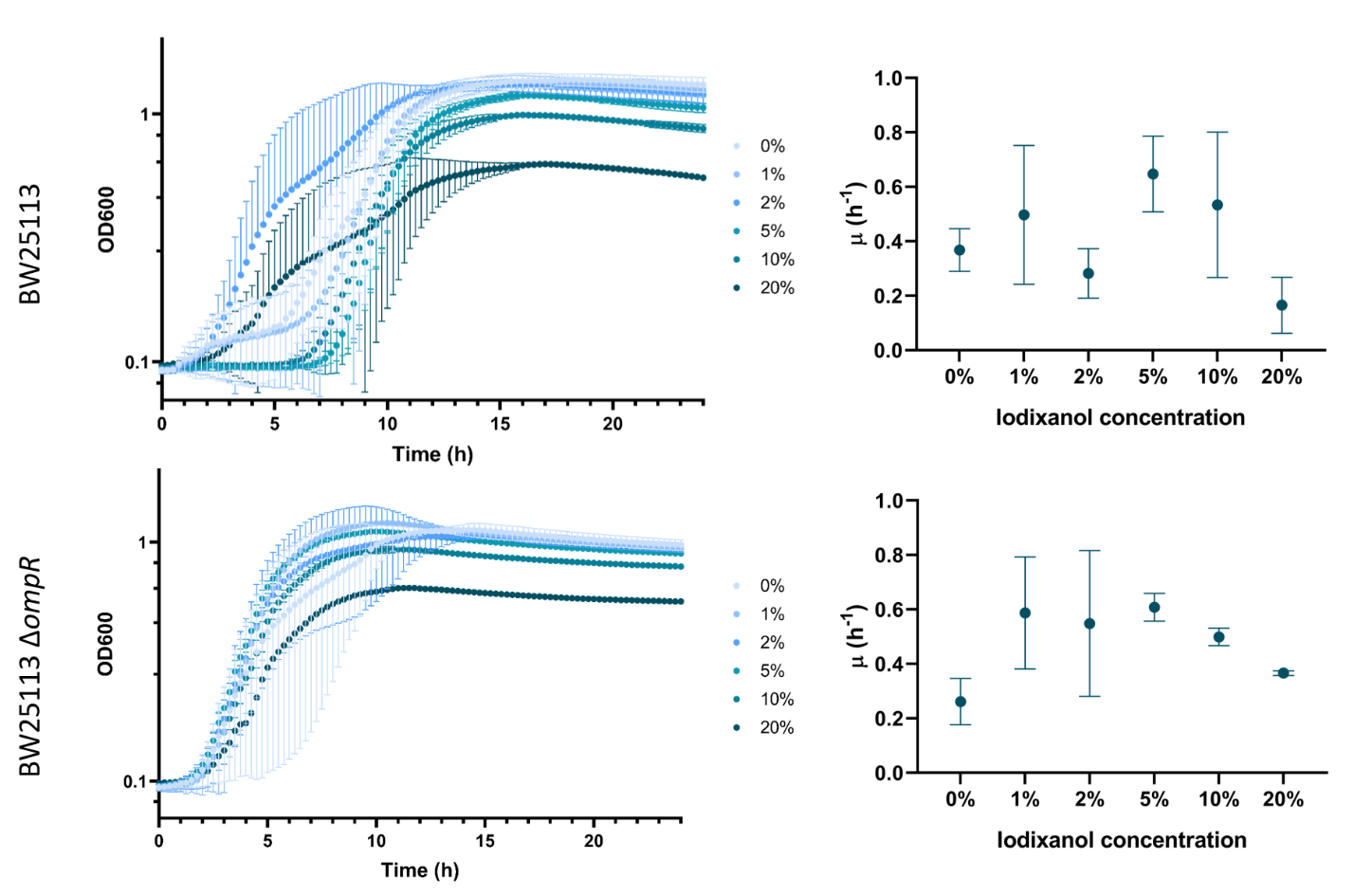

Supplementary Figure 4: Growth curves (left panels) and specific growth rates (right panels) for the parental strain BW25113 and the mutant strain Δ*ompR*, grown in LB broth with increasing v/v concentrations of iodixanol. Growth curves are plotted with the y axis in logarithmic scale. Each data point is the average across 5 repeats, and error bars represent standard deviations.

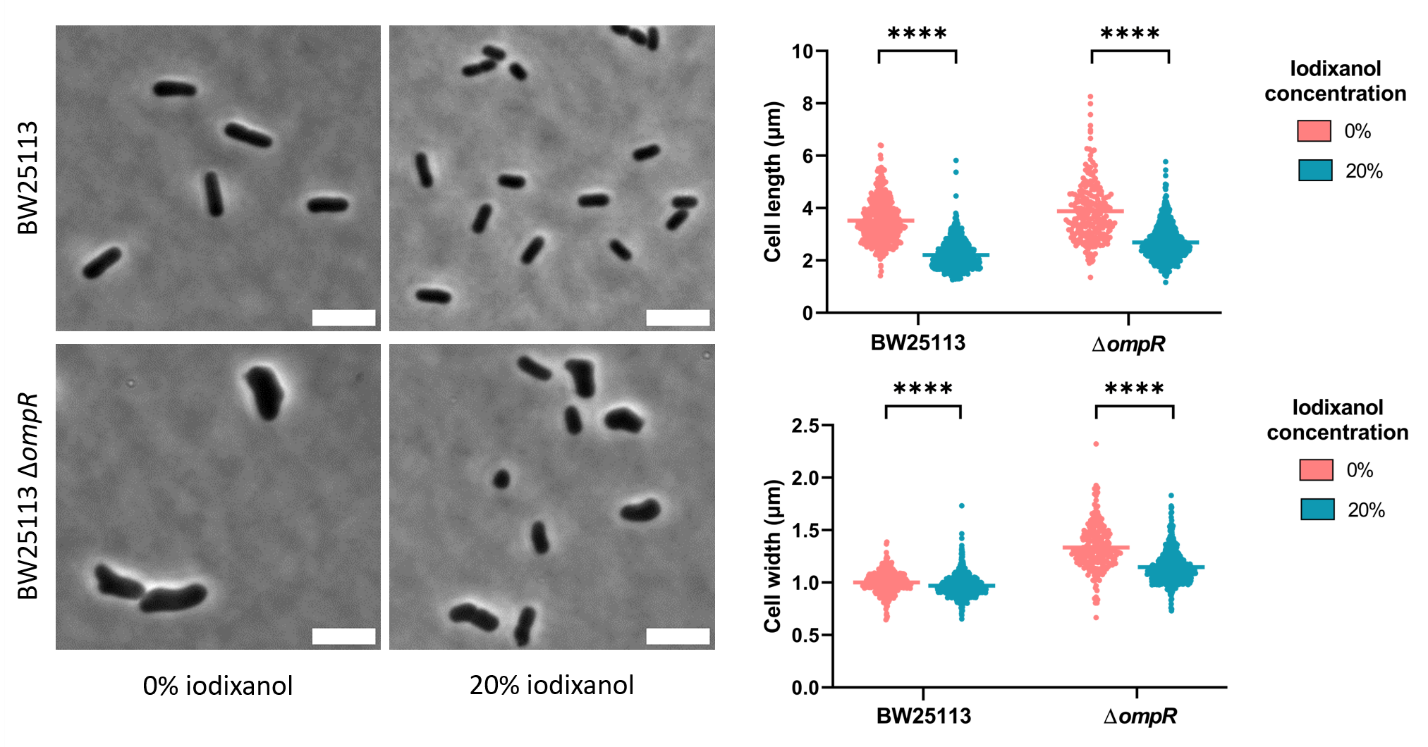

Supplementary Figure 5: Single-cell phenotype of the parental strain BW25113 and of the Δ*ompR* mutant strain, grown in LB broth and in LB broth with 20% iodixanol (v/v). A reduction in cell length of more than 30% can be observed for both BW25113 and Δ*ompR* after growth in medium containing iodixanol (p = 1.32 ⨯ 10^-167^ and p = 6.74 ⨯ 10^-40^ respectively). The Δ*ompR* strain also exhibits a 14% reduction in cell width (p = 1.02 ⨯ 10^-13^), almost five times higher than the parental strain’s 3% reduction (p = 8.57 ⨯ 10^-8^). The number of individual cells analysed in this experiment are n = 444 (BW25113 in LB medium), n = 1197 (BW25113 in LB medium with 20% iodixanol v/v), n = 232 (Δ*ompR* in LB medium) and n = 762 (Δ*ompR* in LB medium with 20% iodixanol v/v). Average values are compared using a Kruskal-Wallis statistical test. Scale bars: 5 µm.
